# Microbial metabolomics’ latest SICRIT: Soft ionization by Chemical Reaction in-Transfer mass spectrometry

**DOI:** 10.1101/2024.07.17.604007

**Authors:** Allyson McAtamney, Allison Ferranti, Denise A. Ludvik, Fitnat H. Yildiz, Mark J. Mandel, Taylor Hayward, Laura M. Sanchez

## Abstract

Microbial metabolomics studies are a common approach to identifying microbial strains that have a capacity to produce new chemistries both *in vitro* and *in situ*. A limitation to applying microbial metabolomics to the discovery of new chemical entities is the rediscovery of known compounds, or “known unknowns.” One contributing factor to this rediscovery is the majority of laboratories use one ionization source–electrospray ionization (ESI)–to conduct metabolomics studies. Although ESI is an efficient, widely adopted ionization method, its widespread use may contribute to the re-identification of known metabolites. Here, we present the use of a dielectric barrier discharge ionization (DBDI) for microbial metabolomics applications through the use of soft ionization chemical reaction in-transfer (SICRIT). Additionally, we compared SICRIT to ESI using two different *Vibrio* species–*Vibrio fischeri,* a symbiotic marine bacterium, and *Vibrio cholerae*, a pathogenic bacterium. Overall, we found that the SICRIT source ionizes a different set of metabolites than ESI, and it has the ability to ionize lipids more efficiently than ESI in positive mode. This work highlights the value of using more than one ionization source for the detection of metabolites.

## Introduction

Microbes have historically been a major source of biologically active compounds.^1^ Bacteria produce and utilize metabolites from both primary metabolism and secondary metabolism, creating what are commonly referred to as natural products (NPs). Natural products have found applications in drug discovery, chemical ecology, and human microbiomes.^2^ The largest impediment to finding new natural products with unique chemical scaffolds has been the high rate of rediscovery of “known unknowns.”^3^ A contributing factor to this rediscovery has been that many labs typically employ a solvent extraction on a microbial culture, and subsequent electrospray ionization-liquid chromatography-tandem mass spectrometry (ESI-LC-MS/MS) analyses.^4^ Although ESI is a widely adopted and powerful method, it may partially contribute to preferentially ionizing the same types of chemical entities.^5^

Dielectric barrier discharge ionization (DBDI) is a newer source for the ionization and subsequent detection of volatile compounds.^6^ The commercially available soft ionization chemical reaction in-transfer (SICRIT) source (Plasmion GmbH, Augsburg, GER) is a DBDI source used at atmospheric pressure, and can be easily coupled onto most instruments’ MS inlet, making it largely vendor agnostic **(Figure S1)**. Recently, SICRIT has been used to characterize compounds in real-time breath samples, murine brain tissue, yak milk, and car exhaust, to name a few.^7–10^ The results of these studies show that the SICRIT source has the ability to ionize volatile compounds and favors smaller, less polar metabolites, slightly overlapping with electron ionization (EI) **(Figure S2)**. Overall, these studies have demonstrated SICRIT’s ability to differentially ionize volatile and nonpolar compounds, and we sought to test whether this source could ionize differential microbial metabolites compared to ESI based ionization of the same extracts.

Symbiotic microbial communities such as host-microbe systems have long been recognized as proficient producers of NPs.^11,12^ The symbiotic relationship between *Vibrio fischeri* and the light organ of the Hawaiian bobtail squid, *Euprymna scolopes*, is a well-studied, robust, host-microbe system consisting of only one host and one microbe.^13–16^ Recent findings from our group, along with prior studies, provide evidence that *V. fischeri* has the ability to produce secondary metabolites that contribute to the symbiosis.^17–21^ *V. fischeri’s* ability to produce bacterial biofilms greatly contributes to its ability to colonize its squid host.^22^ For this reason, we chose to use two different mutants of *V. fischeri*–a strong biofilm producer that overexpresses positive regulator RscS and lacks negative regulator BinK, termed “biofilm up,” and a weak biofilm producer containing an overexpression of the same negative biofilm regulator, termed “biofilm down” **(Table S1)**.^23,24^ We also tested extracts from pathogenic Vibrio species, *V. cholerae,* the causative agent of the disease cholera. Biofilm formation increases environmental survival and infectivity of *V. cholerae.* We used two variants of wild-type *V. cholerae:* “smooth,” which does not produce a wrinkly phenotype on agar, and “rugose,” a variant with enhanced biofilm-forming ability that produces a wrinkly phenotype on agar **(Table S1)**.^25^ The use of two biological systems in this study allows us to ensure our results are robust. To our knowledge, this is the first instance of using DBDI-MS for applications in microbial metabolomics. Additionally, we sought to compare SICRIT to ESI to understand how either source can be used to identify both known and unknown microbial metabolites.

## Materials and Methods

### *V. fischeri* microbial culture

All strains **(Table S1)** were inoculated from frozen stocks into 5 mL of LBS medium with 100 µg/mL kanamycin and incubated at 225 rpm at 25 ℃ overnight. Each overnight culture was normalized to OD_600_ 0.1, and 50 µL of the normalized culture were spread onto a 100 mm LBS-Kanamycin agar plate to create a bacterial lawn. These were grown for 96 hours at 25 ℃.

### *V. cholerae* microbial culture

Both strains **(Table S1)** were streaked onto a 100 mm petri dish with 20 mL of LB agar from frozen stocks and incubated at 30 ℃ overnight.

### Extraction methods

For all microorganisms, cultures were scraped off of agar plates using a metal spatula, avoiding the collection of agar. The scrapped material was placed in scintillation vials with 2 mL MeOH and sonicated for 1 hr. Samples were centrifuged, sterile filtered using a 0.2 micron nylon filter, and dried *in vacuo*. Samples were then re-dissolved in 1:1 MeOH:H_2_O at a concentration of 1 mg/mL and placed in LCMS vials for analysis.

### Chromatography conditions

A 2.1 x 50 mm Agilent Poroshell 120 EC-C18 column with pore size 1.9 µm was used on a Bruker Elute UPLC (Bruker Daltonik, Billerica, MA). The column was equilibrated to 95% A (H_2_O + 0.1% formic acid) and subjected to a 10 minute gradient from 5-100% B (MeOH + 0.1% formic acid) at 0.5 mL/min with an injection volume of 5 µL.

### SICRIT source settings

The SICRIT source was placed onto the MS inlet, and the LC-SICRIT module was installed **(Figure S1)**. The source was heated to 400 ℃ and ran with an amplitude of 1800 V and frequency of 45000 Hz. Dry gas was turned off, and nebulizer gas (N_2_) was run at 2.5 L/min. The nebulizer gas line from the MS was directed to the nebulizing and sheath gas inputs of the module by an 1/8th inch Swagelok tee with a needle valve to control flows. Gas flow was primarily directed to the nebulizer, with the remainder to the sheath, where these gas streams help to nebulize, dry, and transport the LC effluent to the SICRIT source.

### ESI source settings

All data was collected using a Bruker timsTOF FleX (Bruker Daltonik, Billerica, MA). Nebulizer gas (N_2_) was set to 2.8 bar, and dry gas was set to 10.0 L/min. Dry temp was set to 230 ℃.

### MS data acquisition

All samples were run in positive mode from *m/z* 100-2000. LC-MS data acquisition was performed in three technical replicates at a spectra rate of 6 Hz. LC-MS/MS data was collected in triplicate using data-dependent acquisition (DDA) at an MS spectra rate of 10 Hz and MS/MS spectra rate at 16 Hz with the top 9 precursors selected for fragmentation at the collision energies and isolation widths in **Table S2**.

### Statistical analysis

All statistical analyses were performed using Metaboscape (Bruker Daltonik, Billerica, MA). The raw MS^1^ Bruker files were imported using T-ReX^Ⓡ^ 3D. The minimum number of features for extraction and result was 3/18, so a feature must have been present in 3 files to appear in the feature list. The peak intensity threshold was 2500, retention time spanning the gradient only. Ions used for extraction include [M+H]^+^, [M+Na]^+^, [M+NH_4_]^+^, and [M+K]^+^. Modifications included a loss of water [M-H_2_O+H]^+^. Following the generation of a feature list, PCA and Student’s t-test were performed.

### MZmine preprocessing

Bruker data was converted using MSConvert^26^ and uploaded to MZmine v. 2.37.^27^ For MZmine data preprocessing, the Global Natural Products Social molecular networking (GNPS) documentation titled “FBMN with MZmine’’ was followed.^27–31^ The noise level for MS^1^ was set to 2500, while the MS^2^ noise level was set to 300. A mass tolerance of 0.05 Da was used for all preprocessing. The minimum peak height for chromatogram deconvolution was 5000, with a peak duration range of 0.03 min-0.4 min and baseline level of 1000. Once the chromatograms were built, deconvoluted, and deisotoped, the join aligner was used with a RT tolerance of 0.15 min. The remainder of the preprocessing for ion identity molecular networking (IIMN) were performed following the GNPS documentation titled “IIN with MZmine’’ and all settings matched the recommended values from the documentation.^29^

### Molecular networking

Feature-based molecular networking (FBMN) via GNPS was performed using mass tolerances of 0.05 Da, Minimum cosine 0.7, 5 minimum matched fragments, and all remaining default settings. Ion identity molecular networking (IIMN) was also performed using this workflow and settings.^29^ All networks were analyzed in Cytoscape version 3.6.1.^32^ Mirror matches were obtained on GNPS using classical molecular networking with the same settings as FBMN, and they were visualized using the Metabolomics Spectrum Resolver.^33^

### Lipidomics analysis

The raw MS^2^ Bruker files were imported using T-ReX^Ⓡ^ 3D. The minimum number of features for extraction and result was 3/18. The peak intensity threshold was 2500, retention time spanning the gradient only. Ions used for extraction include [M+H]^+^, [M+Na]^+^, [M+NH_4_]^+^, and [M+K]^+^. Modifications included a loss of water [M-H_2_O+H]^+^. MS/MS import was used. Annotations were added in Metaboscape using the target list function. The LIPID MAPS database was used as a target list, along with the Bruker Lipids database.^34^ The annotated feature list was exported, and only annotations validated with MS/MS spectra were used for further analysis. Using the LIPID MAPS database, lipid category and main class were added to the feature list. Bar charts were generated in R using ggplot2.^35^

## Results and Discussion

To assess the differences in ionization potential between the two different sources, we first assessed the number of features detected by each source. Although there were comparable numbers of features detected using both sources, each source differentially ionized a subset of features **(Figure 1A)**. Using the precursor masses from feature lists created in Bruker’s Metaboscape software, we produced box-and-whisker plots showing the distribution of *m/z* values ionized by each source. Both sources ionized features in a similar mass range, with the SICRIT source spanning slightly higher into the mass range **(Figure S2)**. For microorganisms, we can infer that each source ionizes a different set of metabolites with a significant overlap between the two.

**Figure 1.**
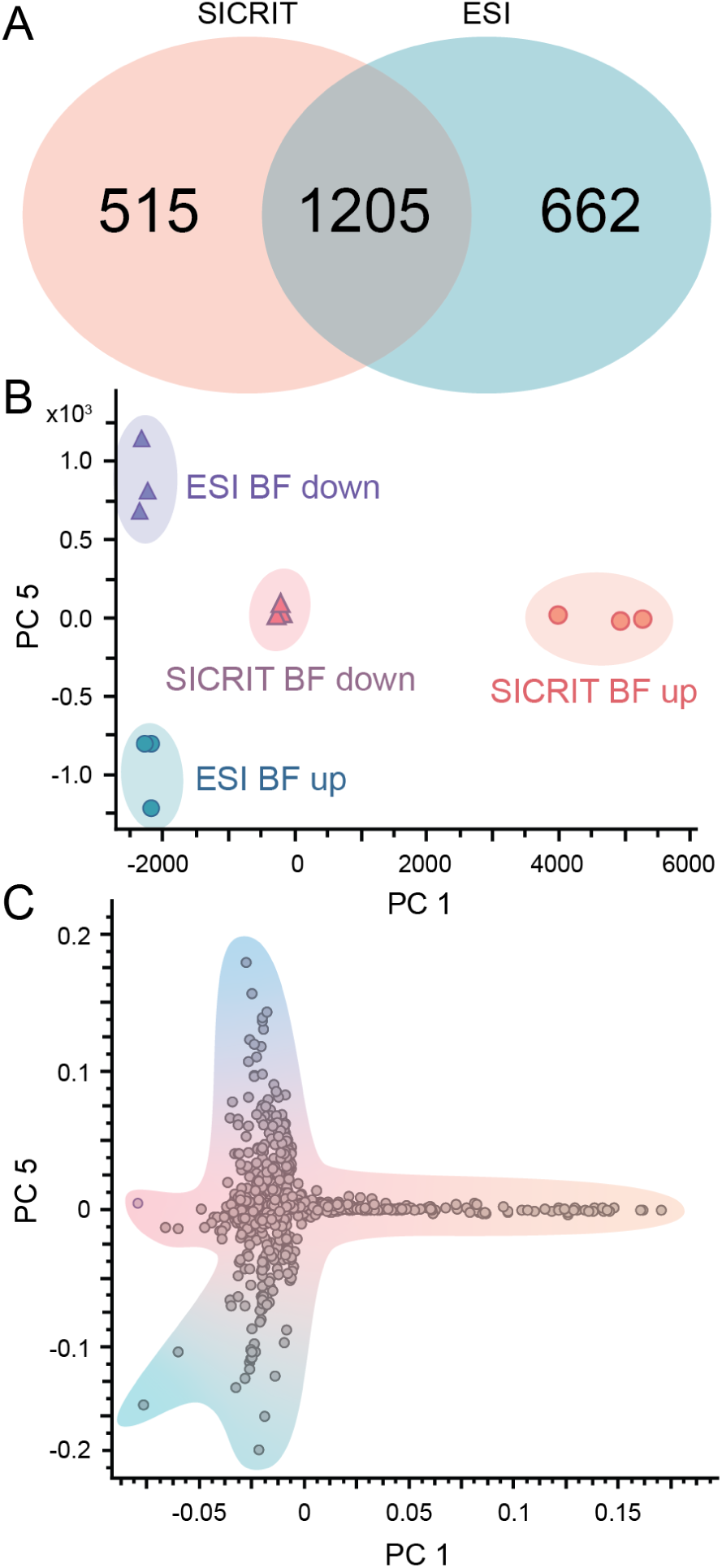
Statistical comparisons between SICRIT and ESI *V. fischeri* data. **A)** Venn diagram with the number of features detected in each source for all biological conditions combined. **B)** PCA 5 vs. PCA 1 scores plot. Each point represents one technical LC-MS/MS replicate. **C)** PCA 5 vs. PCA 1 loadings plot. Each point represents one feature contributing to the overall variance.

Next we sought to assess whether the SICRIT source was able to ionize biologically relevant ions compared to the ESI source. Using principal component analysis (PCA), we found that PC1 and PC5 separated features based on both biological condition (high vs low biofilm) and ionization source **(Figure 1B and Figure S3)**. To identify features contributing to this statistical separation between experiment conditions, we analyzed the loadings plot for PC5 vs. PC1 **(Figure 1C)** and selected features contributing most to the statistical variance. We selected features upregulated in the biofilm-up conditions for both sources and analyzed their extracted ion chromatograms (EICs) to ensure the features were present throughout all technical replicates **(Figure S4 and S5)**. We observed that many of the features contributing to the variance showed very low levels of ionization in the opposing source. These data further suggest that the metabolites ionized by each source are unique and that each source is sensitive enough to identify differences in metabolism by microbial strain based on their underlying biological differences.

Following these analyses, we were interested in better understanding the identities of the features driving the differences between the sources. We used Feature-based molecular networking (FBMN) in the GNPS online environment to further assess the data.^28^ In contrast to classical molecular networking, FBMN allows for the use of ion identity molecular networking (IIMN), which provides information about the type of adducts detected throughout molecular networks.^29^ Using this workflow, we were able to visualize various properties of the *V. fischeri* molecular network, including precursor mass, adduct, biological condition, and molecule class **(Figure 2)**.

**Figure 2.**
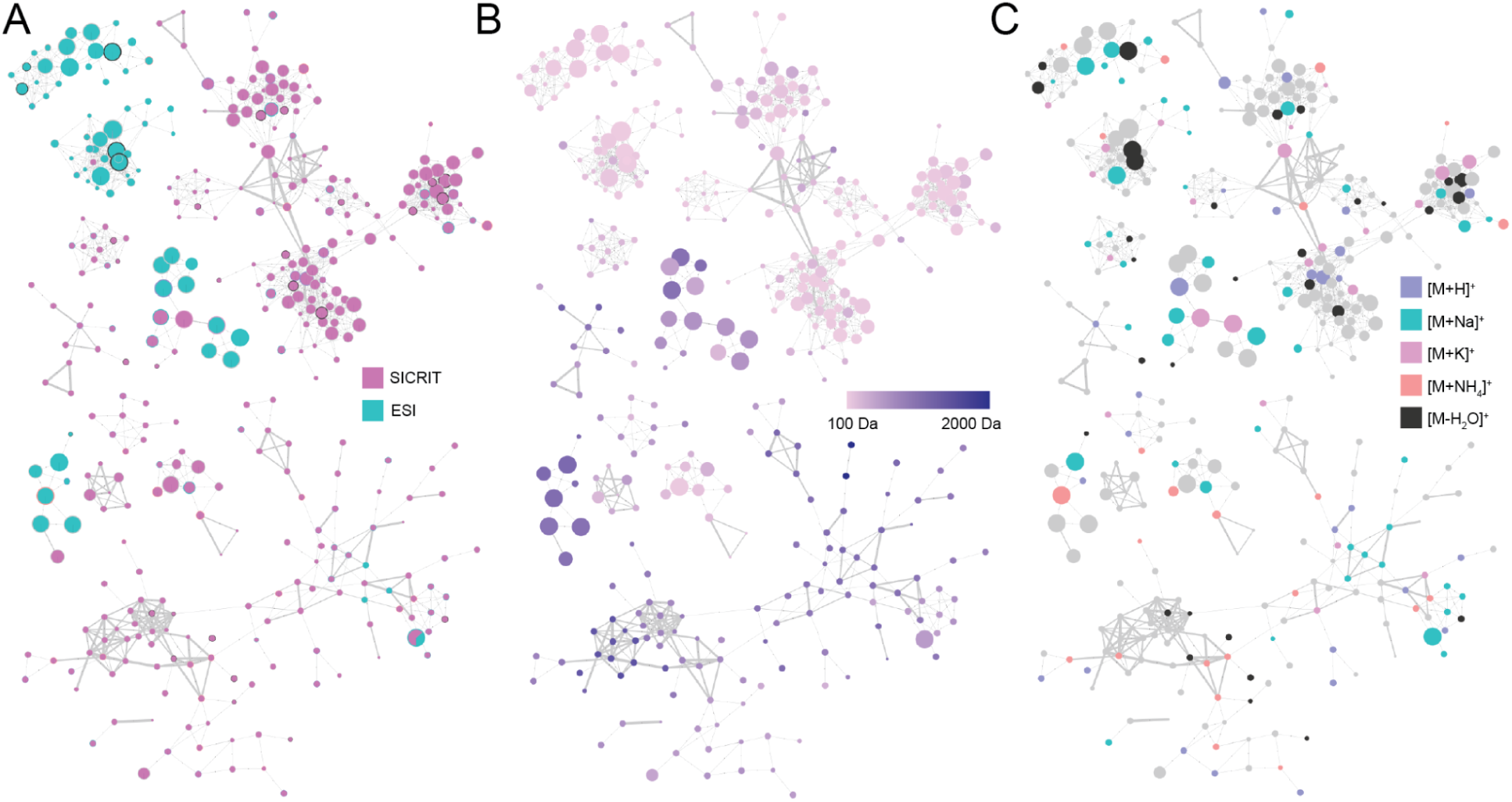
Feature-based molecular networks for *V. fischeri* dataset. Color coded based on **A)** Ionization source, **B)** Adduct predicted through IIMN, and **C)** Precursor mass.

By visually analyzing the *V. fischeri* molecular network, we observed that the features ionized by SICRIT clustered separately from features ionized by ESI **(Figure 2A)**. A small number of clusters contained features from both sources, but a majority clustered separately. Overall, there were 1,209 nodes from the SICRIT source, 381 from ESI, and 14 that were present in both sources. Using IIMN, we annotated the adduct of each feature within the network; however, we did not find there was a trend in the adducts produced by either source **(Figure 2B)**. Several [M+H]^+^ adducts were annotated throughout the network, but they were spread out throughout the entire network, suggesting that the SICRIT source is also a soft ionization source. We also observed that doubly charged adducts were infrequent but evenly distributed throughout the network. Considering the SICRIT source was predicted to ionize smaller, more nonpolar metabolites, we were interested in validating this using the molecular network. Contrary to our initial hypothesis, we did not find that either source favored a certain *m/z* range **(Figure 2C)**. To statistically confirm our observations, we created a violin plot and confirmed the lack of statistical significance using a Mann-Whitney Wilcox U-test **(Figure S2)**. These findings suggest that the SICRIT source is a soft ionization source, and it can ionize microbial metabolites spanning a wide size range. Additionally, our hypothesis is further validated by the differential clustering of features ionized by each source. The full *V. fischeri* molecular network can be found in **Figure S6.**

Next, we were interested in assigning identifications to microbial metabolites within our *V. fischeri* molecular network. We were interested in both metabolites produced by the SICRIT source only, as well as metabolites upregulated in the biofilm-up mutant. We hypothesized that most annotations would be less polar metabolites such as lipids, which we were able to easily detect using the SICRIT source in positive mode. We were able to annotate many lipid classes throughout our network such as ceramides (Cers), diacylglycerols (DAGs), and N-acylethanolamines (NAEs), and a majority of lipids within the molecular networks were detected in the SICRIT source data only. For this study, we decided to focus on phosphoethanolamines (PEs) due to their importance in antibiotic resistance in gram-negative bacteria.^36,37^ Several lipid annotations we observed had some of the highest cosine scores throughout the entire network. Although these compounds were detected in higher abundance in the biofilm-down mutant, we observed their presence in both strains, using 1-palmitoyl-2-hydroxy-sn-glycero-3-phosphoethanolamine (PE) for example **(Figure 3A)**.

**Figure 3.**
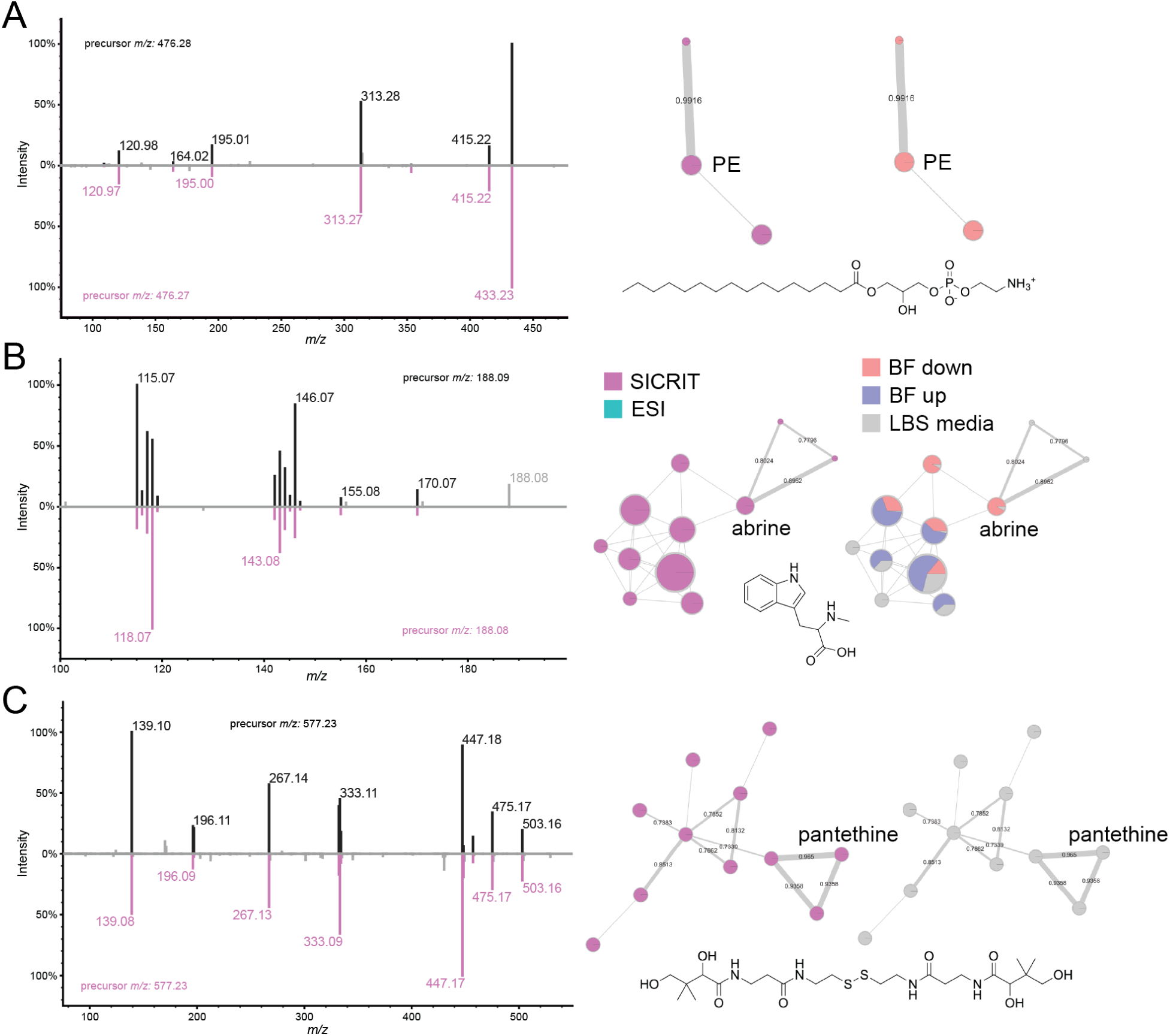
Compounds identified in *V. fischeri* molecular networks. Each compound is shown with its GNPS mirror match between the experimental data (top) and database spectra (bottom).^33^ Next to each mirror match is the molecular network color-coded based on both ionization source (left) and biological condition (right). This data is shown for **A)** abrine, **B)** pantethine, and **C)** 1-palmitoyl-2-hydroxy-sn-glycero-3-phosphoethanolamine (PE)

Despite SICRIT’s strong ability to ionize lipids and other nonpolar compounds, we were able to detect several polar metabolites using the SICRIT source. These include abrine, or *N*-methyl tryptophan, which is a known microbial metabolite **(Figure 3B)**, as well as pantethine, an intermediate in coenzyme A biosynthesis **(Figure 3C)**. Both of these compounds were only detected in the SICRIT source, showing SICRIT’s ability to ionize various types of compounds. A majority of compounds ionized via the SICRIT source that could not be identified via GNPS were below 200 Da, and therefore did not produce many fragments. Considering these metabolites are small and did not ionize via ESI, we hypothesize these metabolites may be more volatile, similar to pheromones. Because of their small size, they remain very difficult to isolate and characterize.

Next, we wanted to validate the findings from *V. fischeri* extracts using a different biological system. To do this, we chose *V. cholerae*, a pathogenic member of the *Vibrionaceae* family. We found that the molecular networks between these two organisms had very similar trends. The SICRIT and ESI features clustered separately once again, and we did not find that either source favored any specific adduct formation **(Figure 4A, 4B)**. We also found both sources to ionize a wide range of metabolite sizes **(Figure 4C)**. Additionally, similar lipids were detected in *V. cholerae*, specifically PEs **(Figure S7)**. The full *V. cholerae* molecular network can be found in **Figure S8.**

**Figure 4.**
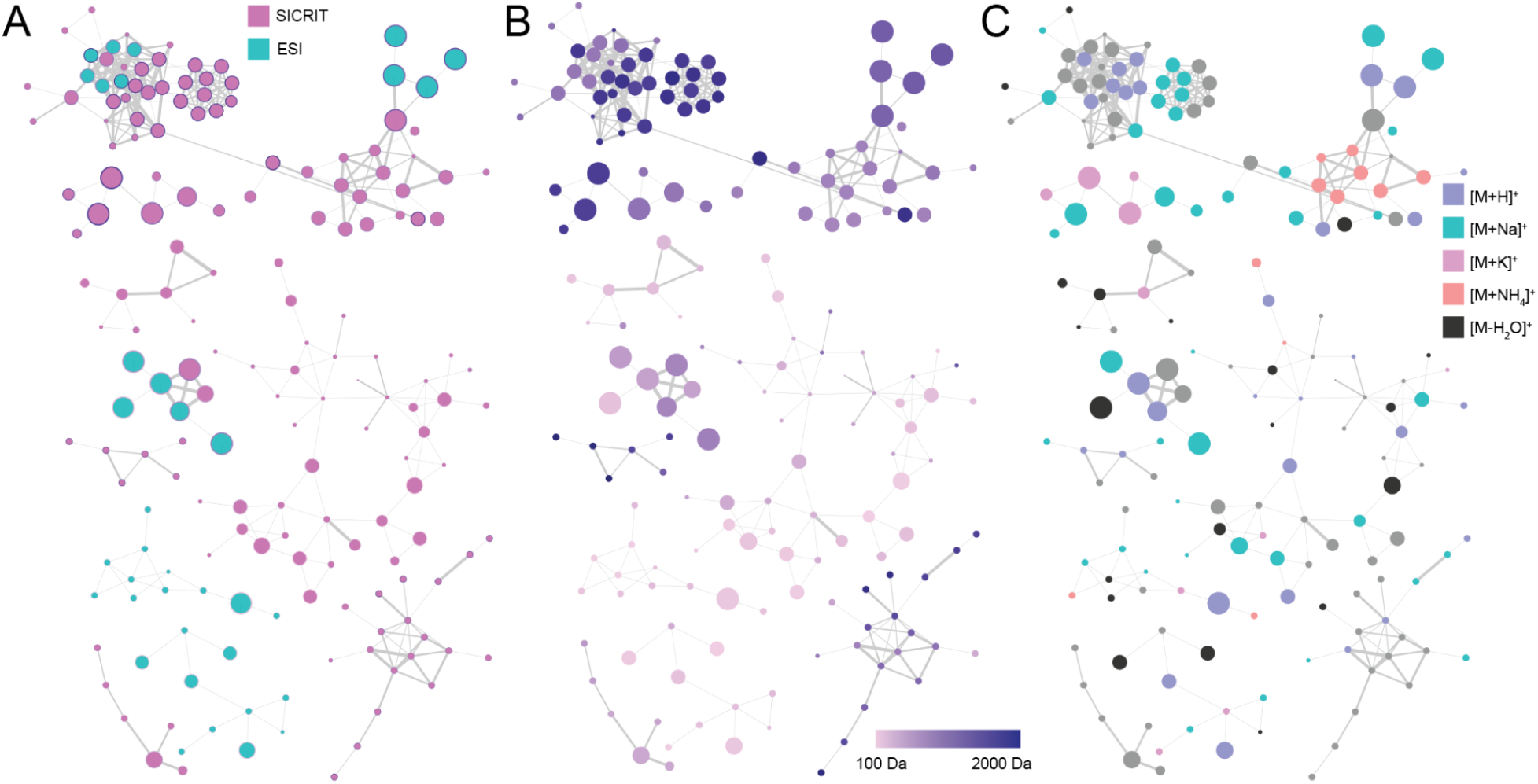
Feature-based molecular networks for *V. cholerae* dataset. Color coded based on **A)** Ionization source, **B)** Adduct predicted through IIMN, and **C)** Precursor mass.

Finally, previous studies have shown the SICRIT source efficiently ionizes lipids in mammalian biological systems.^38^ However, we were interested in understanding how this translates to a microbial system. From our MS^2^-level annotated feature list, we saw that our annotations remained robust in both biological systems, with a majority of annotations belonging to the fatty acyls (FA), sphingolipids (SP), glycerolipids (GL), and glycerophospholipids (GP) LIPID MAPS categories **(Figure 5A)**.^34^ Considering FAs were the most abundant category, we were interested in understanding whether either source favors specific main classes within this lipid category. Using the same *V. fischeri* dataset, we observed that the SICRIT source ionized all annotated FA main classes more efficiently than the ESI source except fatty esters **(Figure 5B)**.

**Figure 5.**
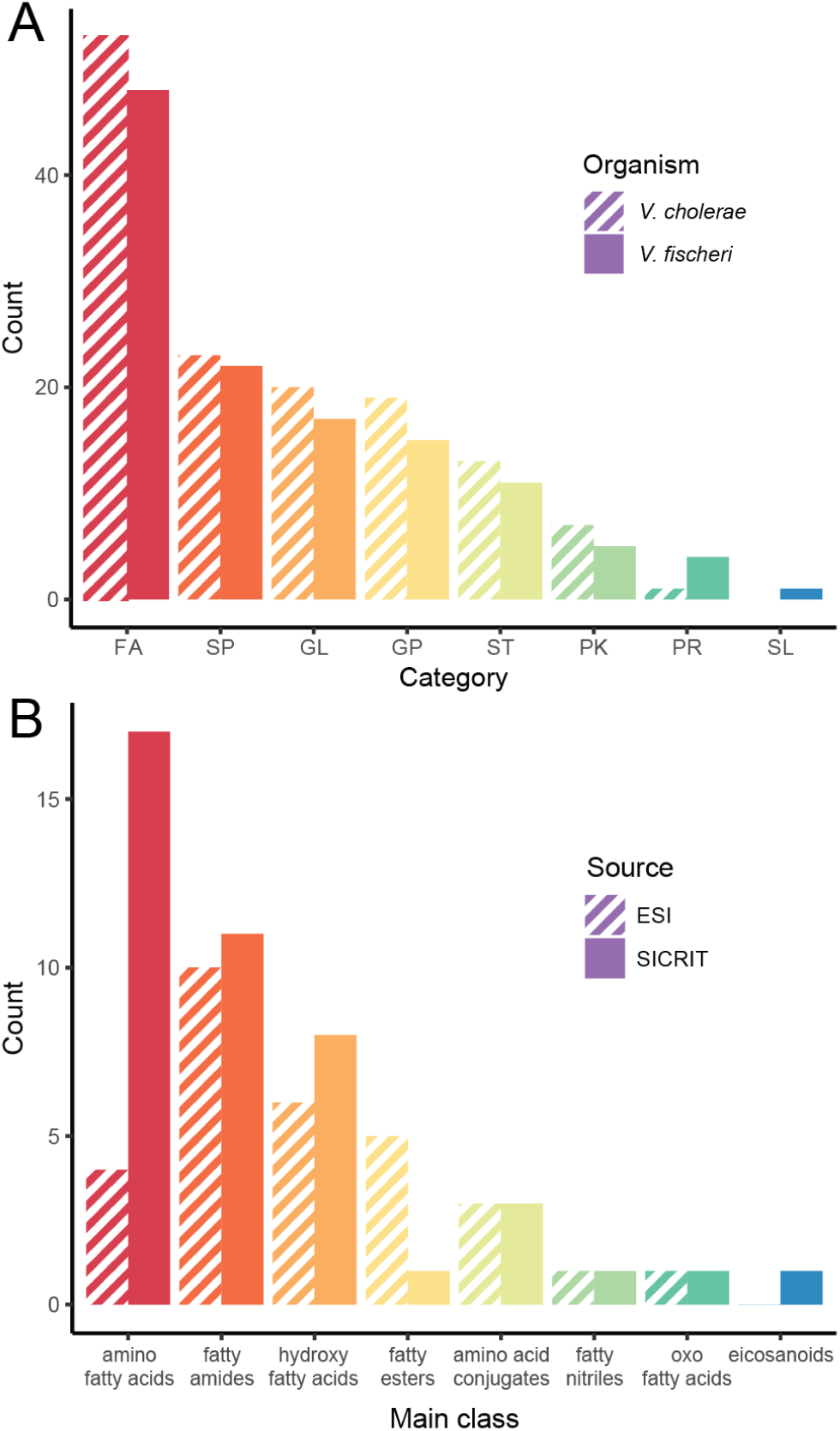
Bar chart of the total number of lipid annotations by **A)** LIPID MAPS category: Fatty Acyls (FA), Sphingolipids (SP), Glycerolipids (GL), Glycerophospholipids (GP), Sterol Lipids (SL), Polyketides (PK), Prenol Lipids (PR), and Saccharolipids (SL).^34^ The striped bars correspond to *V. cholerae* counts, and the solid bars correspond to the *V. fischeri* counts. **B)** *V. fischeri* fatty acyl main class annotation counts by ionization source. The striped bars correspond to ESI annotation counts, and the solid bars correspond to SICRIT annotation counts. All annotations are MS^2^ level and were obtained from Bruker’s Metaboscape software.

## Conclusions

Our findings suggest that the SICRIT ionization source can ionize a different set of microbial metabolites than the conventionally used ESI source. Contrary to our initial hypothesis, the SICRIT source can ionize a wide range of microbial metabolite sizes and produces similar adducts as ESI. It also has the ability to detect polar metabolites, such as pantethine and abrine **(Figure 3B, 3C)**. SICRIT’s increased ability to ionize lipids in positive mode makes it a good candidate for lipidomics studies as compared to ESI, but this should be validated using more targeted lipidomics studies. It is worth nothing that headspace analyses using the SICRIT source from microbes grown directly on agar in Petri dishes resulted in the selective ionization of plasticizers from the Petri dish itself. Overall, there are a number of shared metabolites that can be ionized by both SICRIT and ESI, but SICRIT has the unique ability to ionize a variety of microbial lipids in positive mode beyond that observed using ESI. The features detected in both sources are biologically meaningful following trends in biofilm production.

## Data availability

All raw data is available on the MassIVE database under MSV000095362. *V. fischeri* GNPS job ID:246f05b912f54a96948c12ae41dca13f and *V. cholerae* GNPS job ID:70d1651bf07541dba9f1132b0561e781.

## Acknowledgements

This work was supported by the National Institute of Allergy and Infectious Diseases of the NIH under Award Number R01AI102584 (F.H.Y.), the National Institute of General Medical Sciences of the NIH award R35GM148385 (M.J.M.), and by National Science Foundation grants IOS-2220510 (L.M.S.) and IOS-2220511 (M.J.M.), UCSC Startup funds (L.M.S.).

## Notes

The authors declare the following competing financial interest(s): Taylor Hayward and Allison Ferranti are and were employees of Plasmion GmbH, respectively.

