## Supplemental Information for "Microbial metabolomics’ latest SICRIT: Soft ionization by Chemical Reaction in-Transfer mass spectrometry"

| | | |
|---|---|---|
| **Figure S1.** | LC-SICRIT apparatus………………………………………………………… | S2 |
| **Figure S2.** | Violin plot for precursor masses detected in ESI and SICRIT……………….. | S3 |
| **Figure S3.** | Principal component analysis data for PCAs 1-5…………………………….. | S4 |
| **Figure S4.** | EICs of upregulated features in SICRIT + Biofilm up……………………….. | S5 |
| **Figure S5.** | EICs of upregulated features in ESI + Biofilm up……………………………. | S6 |
| **Figure S6.** | Full *V. fischeri* molecular network…………………………………………… | S7 |
| **Figure S7.** | Mirror matches of phosphoethanolamines in *V. cholerae* data……………….. | S8 |
| **Figure S8.** | Full *V. cholerae* molecular network…………………………………………... | S9 |

**Materials and Methods:**

| | | |
|---|---|---|
| **Table S1.** | Box-and-whisker plot for precursor masses detected in ESI and SICRIT…… | S10 |
| **Table S2.** | Principal component analysis data for PCAs 1-5…………………………….. | S10 |
| **References** | ……………………………………………………………………………... | S11 |

**Figure S1.** Labeled LC-SICRIT apparatus.

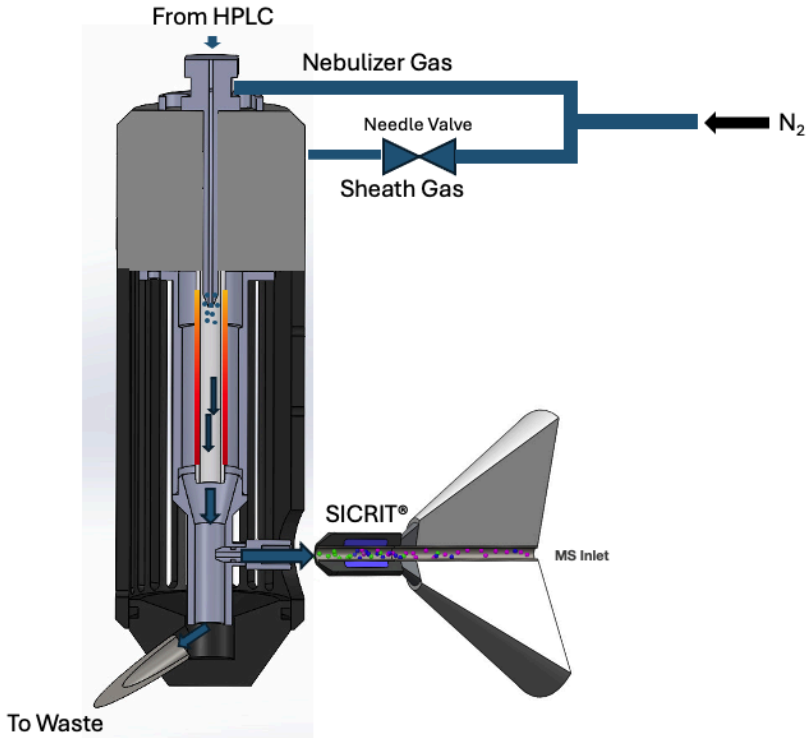

**Figure S2.** Violin plot for precursor masses of features detected in the SICRIT or ESI source. The bold line represents the medium *m/z* that were detected. The mass range for this experiment was 100-2000 Da.

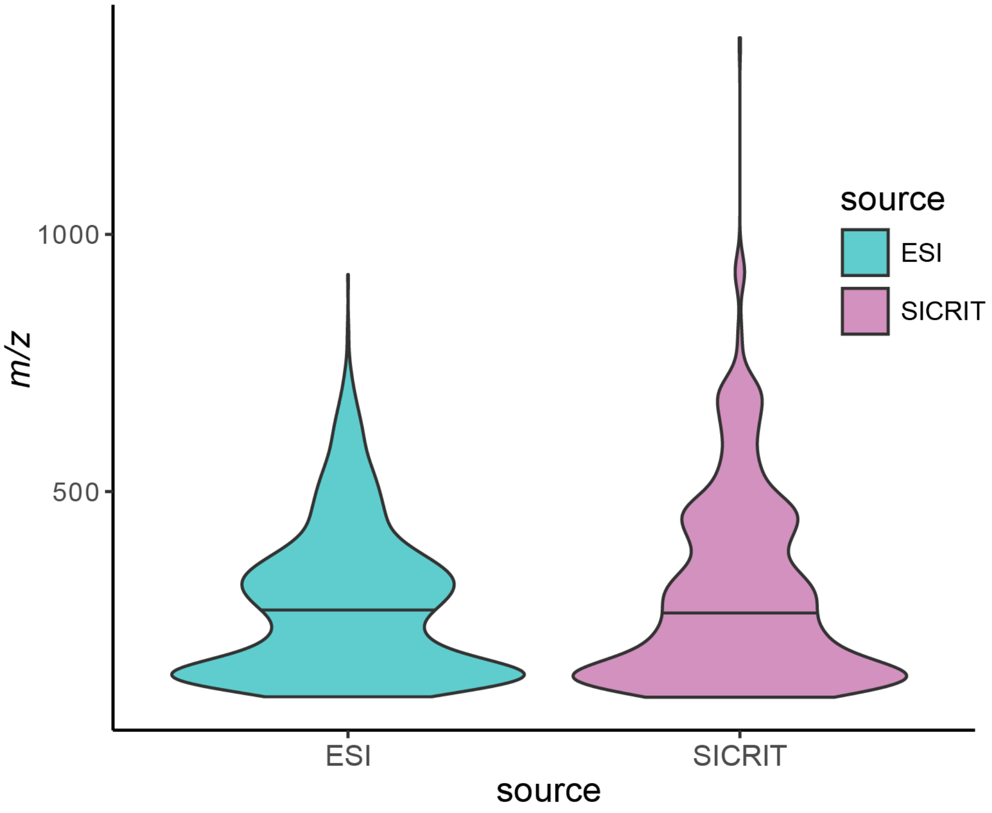

**Figure S3.** Principal component analysis for PCAs 1-5. The intersection of each principal component represents the PCA plot for those components. All x-axes at the bottom represent the plots above. All y-axes on the left represent plots to the right. Each condition contains three technical replicates, each represented by either a circle (biofilm up) or triangle (biofilm down). All ESI replicates are shown in turquoise. All SICRIT replicates are shown in pink.

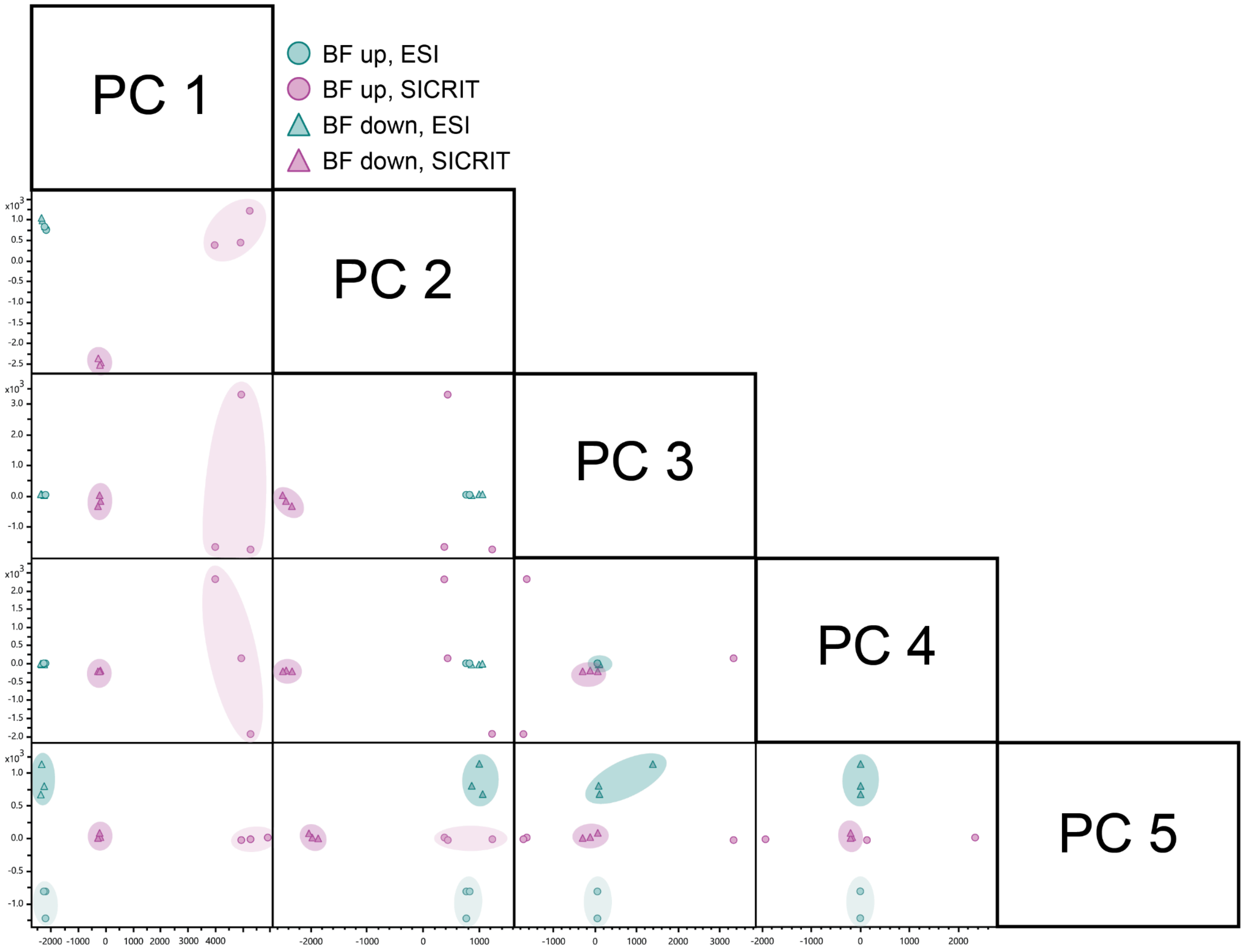

**Figure S4.** Upregulated features detected in the biofilm up condition and ionized by the SICRIT source. Both were identified using the PCA loadings plot. **A)** Extracted ion chromatogram for *m/z* 254.20. **B)** Extracted ion chromatogram for *m/z* 298.31.

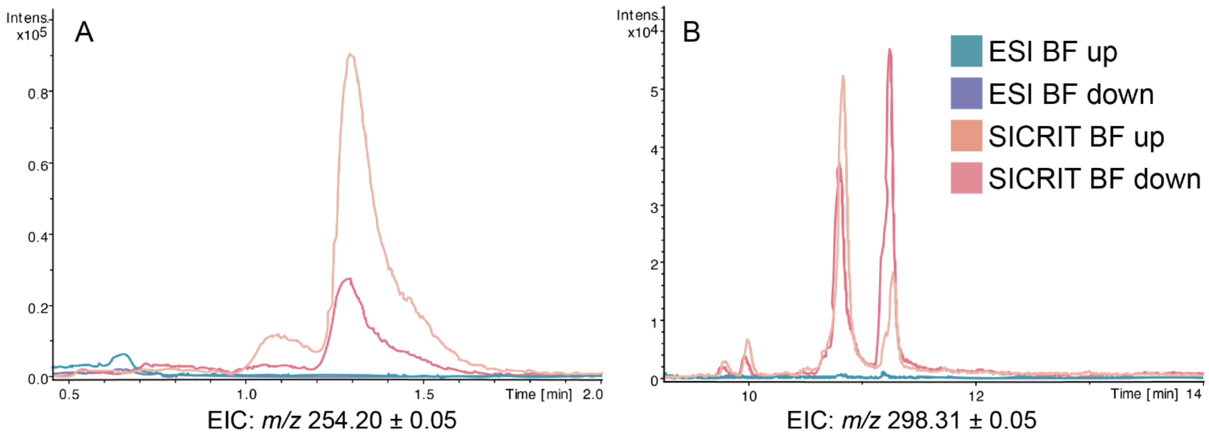

**Figure S5.** Upregulated features detected in the biofilm up condition and ionized by the ESI source. Both were identified using the PCA loadings plot. **A)** Extracted ion chromatogram for *m/z* 381.30. **B)** Extracted ion chromatogram for *m/z* 685.52.

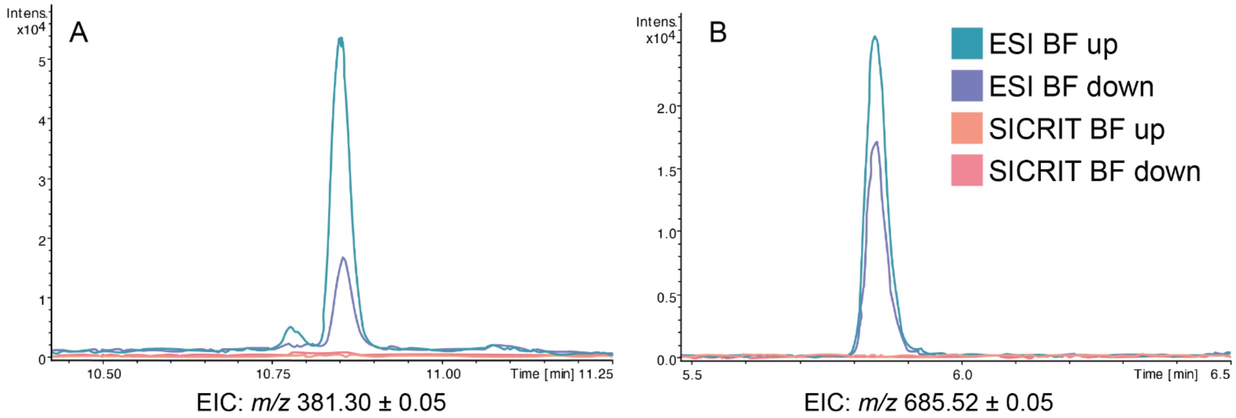

**Figure S6.** Mirror match spectra for phosphoethanolamines identified in *V. cholerae* extracts. Top = experimental MS/MS spectrum. Bottom = GNPS library MS/MS spectrum. Both annotations were present in both smooth and rugose variants. **A)** Mirror match for 1-palmitoyl-2-hydroxy-sn-glycero-3-phosphoethanolamine. **B)** Mirror match for 1,2-di-(9Z-octadecenoyl)-sn-glycero-3-phosphoethanolamine.

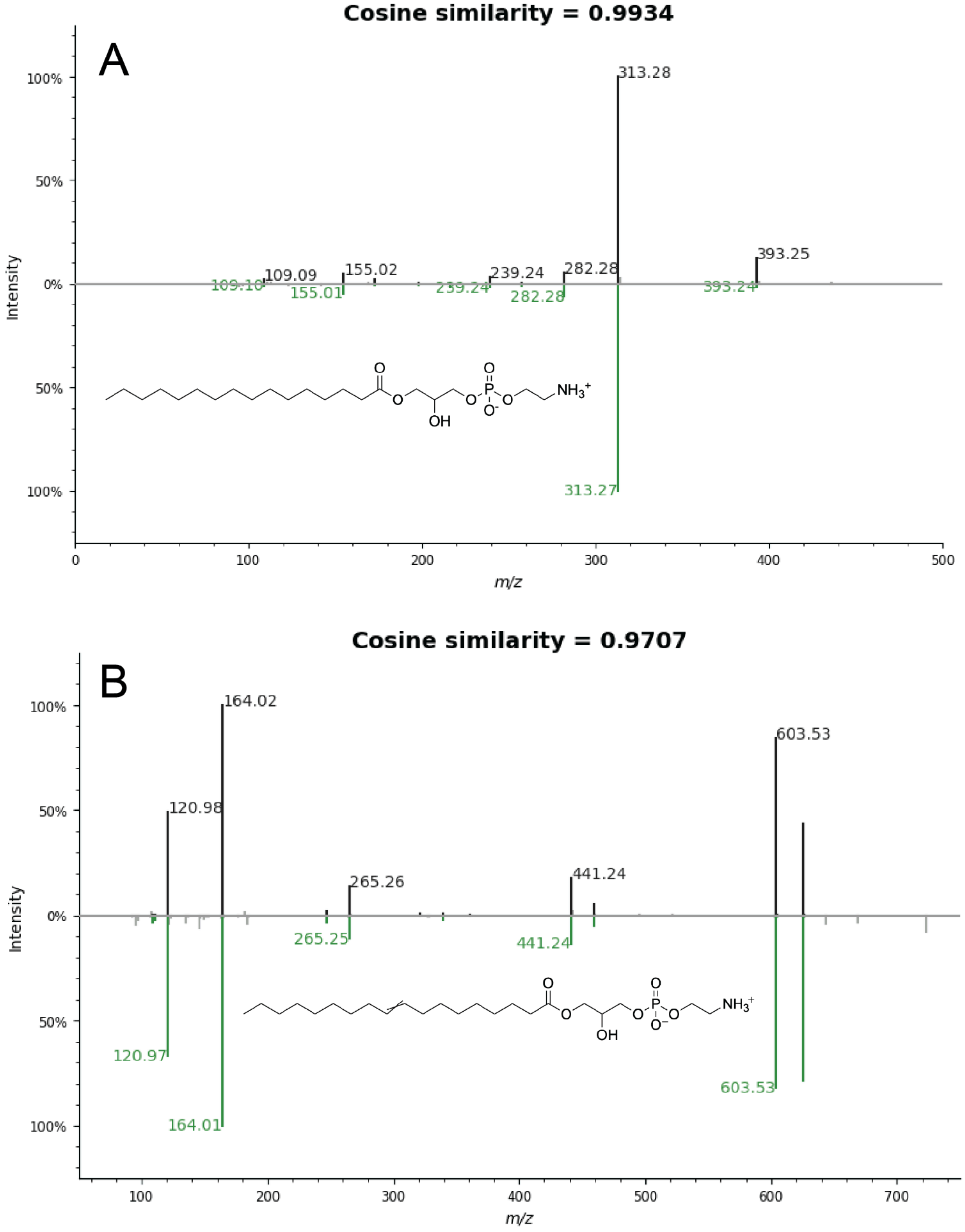

**Figure S7.** Full *V. fischeri* molecular network, color coded by ionization source. All SICRIT nodes are shown in pink, and all ESI nodes are shown in turquoise.

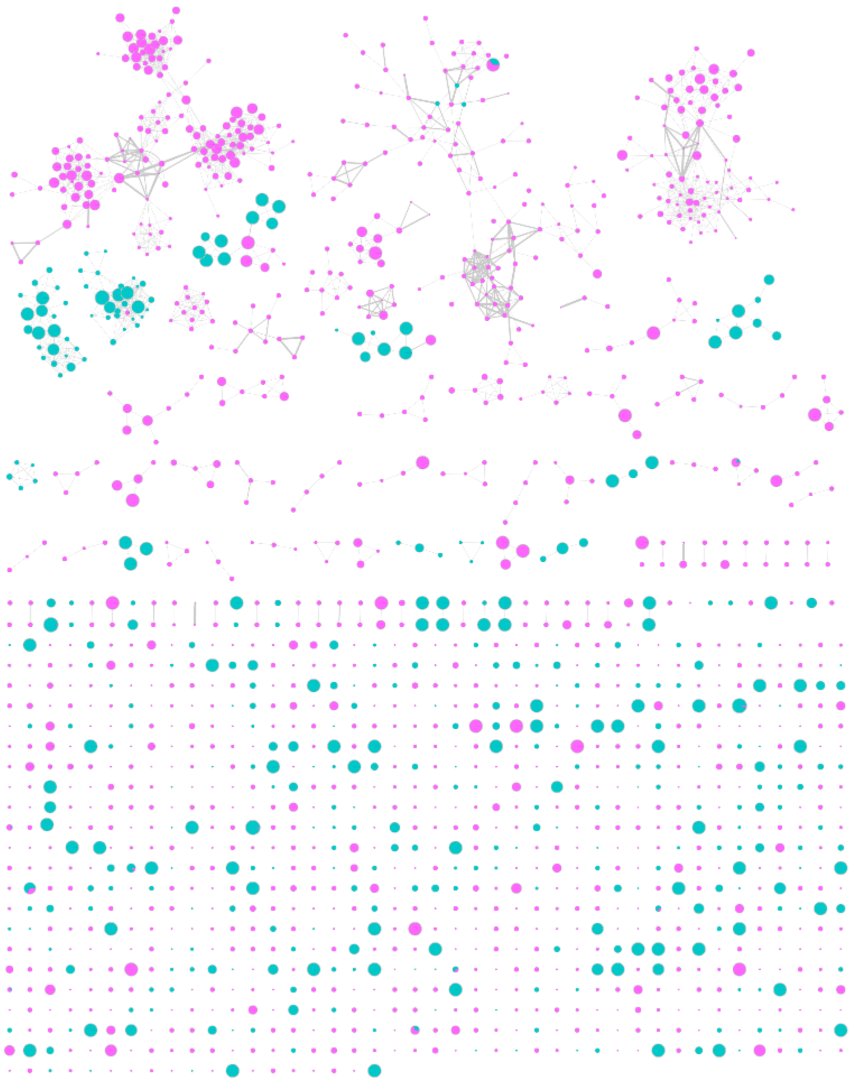

**Figure S8.** Full *V. cholerae* molecular network, color coded by ionization source. All SICRIT nodes are shown in pink, and all ESI nodes are shown in turquoise.

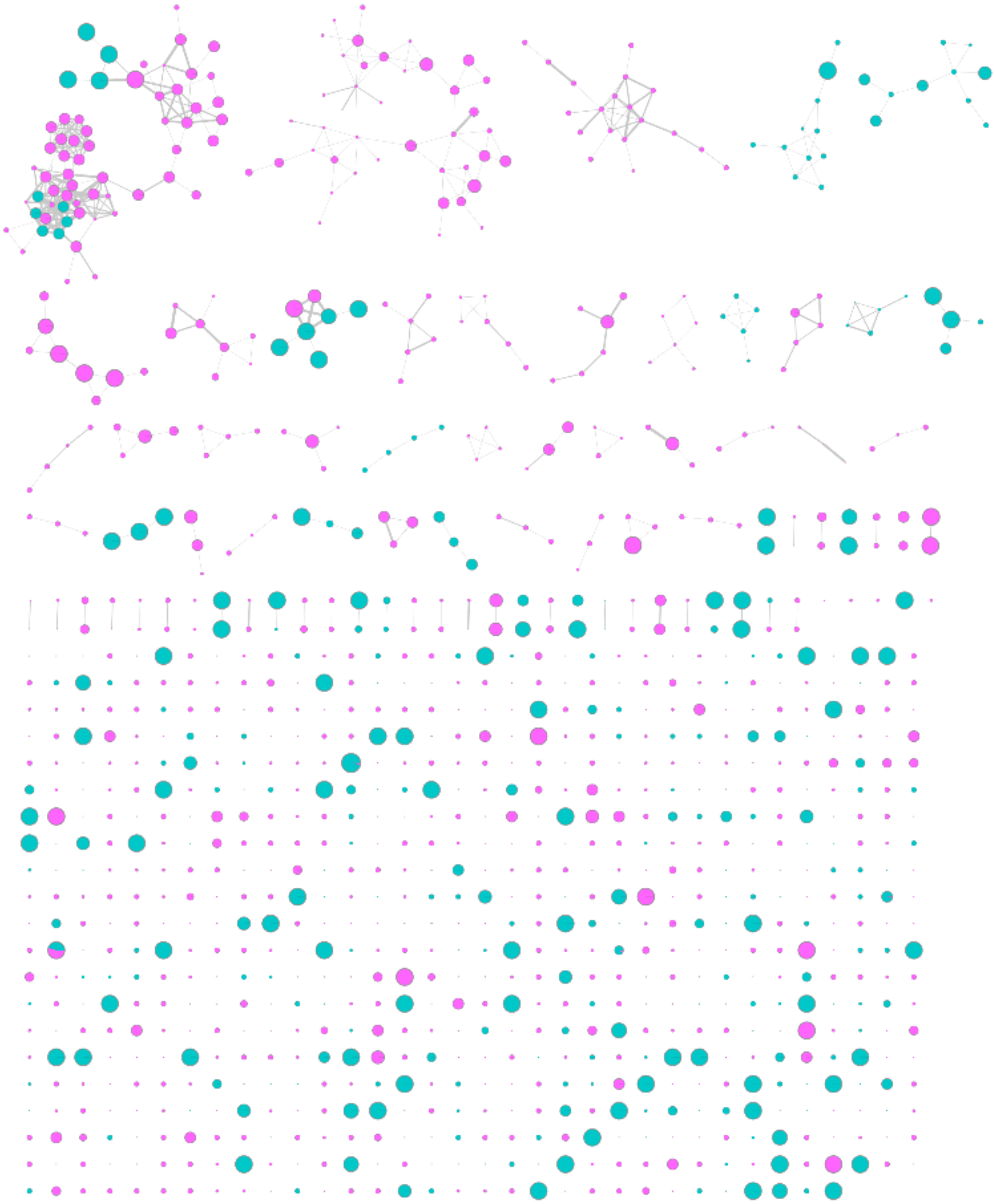

**Materials and Methods**

**Table S1.** Strain table for all organisms used in this study.

| **Organism** | **Common term** | **Strain** | **Genotype** | **Citation** |
|---|---|---|---|---|
| *V. fischeri* | Biofilm down | MJM2386 | ES114 pBinK | (*1*) |
| *V. fischeri* | Biofilm up | MJM2389 | ES114 *rscS** Δ*binK* pVSV104 | (*2*) |
| *V. cholerae* | Smooth | Fy_Vc_1 | O1 El Tor A1552, wild type, Rifr | (*3*) |
| *V. cholerae* | Rugose | Fy_Vc_2 | O1 El Tor A1552, rugose variant, Rifr | (*3*) |

**Table S2.** Collision energy table for MS/MS acquisition.

| **Mass (m/z)** | **Isolation width (m/z)** | **Collision Energy (eV)** |
|---|---|---|
| 100 | 1.00 | 25 |
| 500 | 2.00 | 35 |
| 1000 | 3.00 | 40 |
| 1300 | 4.00 | 55 |

**References.**


1. Rotman Ella R., Bultman Katherine M., Brooks John F., Gyllborg Mattias C., Burgos Hector L., Wollenberg Michael S., and Mandel Mark J. (2019) Natural Strain Variation Reveals Diverse Biofilm Regulation in Squid-Colonizing Vibrio fischeri, *J. Bacteriol.*, American Society for Microbiology *201*, 10.1128/jb.00033–19.
2. Brooks John F., and Mandel Mark J. (2016) The Histidine Kinase BinK Is a Negative Regulator of Biofilm Formation and Squid Colonization, *J. Bacteriol.*, American Society for Microbiology *198*, 2596–2607.
3. Beyhan, S., and Yildiz, F. H. (2007) Smooth to rugose phase variation in Vibrio cholerae can be mediated by a single nucleotide change that targets c-di-GMP signalling pathway, *Mol. Microbiol. 63*, 995–1007.
